# Differential chondrogenic differentiation between iPSC-derived from healthy and OA cartilage is associated with changes in epigenetic regulation and metabolic transcriptomic signatures

**DOI:** 10.1101/2022.10.14.512213

**Authors:** Nazir M. Khan, Martha Elena Diaz-Hernandez, Samir Chihab, Priyanka Priyadarshani, Pallavi Bhattaram, Luke J. Mortensen, Rosa M Guzzo, Hicham Drissi

**Author notes:** **Address for correspondence**: Hicham Drissi, Ph.D. Director, Emory Musculoskeletal Research Centre, Professor and Vice Chair Research, Department of Orthopaedics, Emory University School of Medicine, Atlanta, GA-30033, USA.

## Abstract

Induced pluripotent stem cells (iPSCs) are potential cell sources for regenerative medicine. The iPSCs exhibit a preference for lineage differentiation to the donor cell type indicating the existence of memory of origin. Although the intrinsic effect of the donor cell type on differentiation of iPSCs is well recognized, whether disease-specific factors of donor cells influence the differentiation capacity of iPSC remains unknown. Using viral based reprogramming, we demonstrated the generation of iPSCs from chondrocytes isolated from healthy (AC-iPSCs) and osteoarthritis cartilage (OA-iPSCs). These reprogrammed cells acquired markers of pluripotency and differentiated into uncommitted-mesenchymal progenitors. Interestingly, AC-iPSCs exhibited enhanced chondrogenic potential as compared OA-iPSCs and showed increased expression of chondrogenic genes. Pan-transcriptome analysis showed that chondrocytes derived from AC-iPSCs were enriched in molecular pathways related to energy metabolism and epigenetic regulation, together with distinct expression signature that distinguishes them from OA-iPSCs. The molecular tracing data demonstrated that epigenetic and metabolic marks were imprint of original cell sources from healthy and OA-chondrocytes. Our results suggest that the epigenetic and metabolic memory of disease may predispose OA-iPSCs for their reduced chondrogenic differentiation and thus regulation at epigenetic and metabolic level may be an effective strategy for controlling the chondrogenic potential of iPSCs.

## INTRODUCTION

Osteoarthritis (OA) is an inflammatory joint disease in which catabolic cascade of events results in cartilage destruction leading to severe joint pain.^1^ While non-surgical procedures such as NSAID and steroid injections are helpful, the majority of OA cases ultimately undergo joint replacement therapy. The induced pluripotent stem cells (iPSCs) were recently proposed as a promising source to repair cartilage damage.^2;3^ While iPSCs are seriously considered as potential cell sources for regenerative medicine, accumulating evidence suggests that iPSCs from different cell sources have distinct molecular and functional properties.^4–8^ It has been reported that iPSCs derived from various somatic cell types exhibited a preference for differentiation into their original cell lineages.^4;9^ Therefore, the effects of the cellular origin of iPSCs on their lineage-specific differentiation capacity is an important consideration for cell replacement therapies, drug screening, or disease modeling.

Several studies have determined that iPSCs retain a memory of their cellular origin due to residual DNA methylation and histone modification patterns at lineage specific genes. Thus, this residual ‘epigenetic memory’ has been shown to bias their subsequent differentiation into their parental/donor cell lineage.^10–12^ Although it is known that cellular origin of iPSCs influences their differentiation capacity, the contribution of disease-specific factors on the capacity of iPSC for chondrogenic differentiation remains unknown. Examining potential differences between cells that reside in healthy versus OA environments, would provide unique insight into the chondrogenic potential of these cells, and their utility in disease modeling. Since OA articular chondrocytes exhibit different features from healthy articular chondrocytes, we posit that the iPSCs derived from these cell states represent the feature of their physiological origin. Thus, the memory of the cells is not only specific to the tissue of origin but also to the physiological status which further influences the differentiation capacity and ultimately the efficiency of tissue regeneration.

In the present study, we aimed to determine whether iPSCs derived from healthy and diseased (OA) cartilage possess differential chondrogenic potential, and whether OA disease status significantly limits their differentiation capacity. To this end, we derived iPSCs from healthy (AC-iPSCs) and OA chondrocytes (OA-iPSCs) and compared their differentiation capacity into chondroprogenitors and chondrocytes. During differentiation of iPSCs into chondrocytes, we determined the epigenetic and metabolic marks of cellular memory. Our results showed that iPSCs derived from healthy chondrocytes (AC-iPSCs) exhibited an enhanced potential for chondrocyte differentiation as compared to OA-iPSCs. Our data further demonstrate that although reprogramming of OA chondrocytes induced pluripotency, the OA-iPSCs retained the epigenetic and metabolic marks associated with pathological conditions of diseased chondrocytes and retention of this cellular memory influence their chondrogenic commitment and thus regenerative capacity for the cartilage repair. Our findings indicate that regulating the epigenetic modifiers and energy metabolism may be an effective strategy for enhancing the chondrogenic potential of iPSCs derived from chondrocytes.

## RESULTS

### Characterization of iPSCs generated from healthy and OA articular chondrocytes

We previously reported the generation of iPSCs from healthy articular chondrocytes (AC-iPSCs) and performed molecular, cytochemical, and cytogenic analyses to determine the pluripotency of generated iPSCs.^13^ In the present study, we used multiple clones of the previously generated AC-iPSCs (clones #7, #14, and # 15), and compared their pluripotency, progenitor properties and chondrogenic potential to that of newly generated OA-derived iPSCs (OA-iPSCs) (clones #2, #5 and #8) (**Fig. 1A**). These colonies showed positive alkaline phosphatase (ALP) staining, indicating an undifferentiated pluripotent stem cell phenotype of both AC-iPSC and OA-iPSC clones (**Fig. 1B**). Stemness characteristics of these iPSC clones was evaluated via qPCR assessment of key pluripotency marker genes. The mRNA copy number of *SOX2, OCT4, NANOG* and *KLF4* were comparable in AC-iPSCs and OA-iPSCs (**Fig. 1C**) indicating a similar level of stemness identity between these iPSCs. Interestingly, *KLF4* expression was low as compared to the other pluripotency gene in both iPSCs (**Fig. 1C**). Pluripotency was also colonies from both AC-iPSCs and OA-iPSCs showed positive expression of SOX2 and TRA-1-60 proteins (**Fig.1D**).

**Figure 1:**
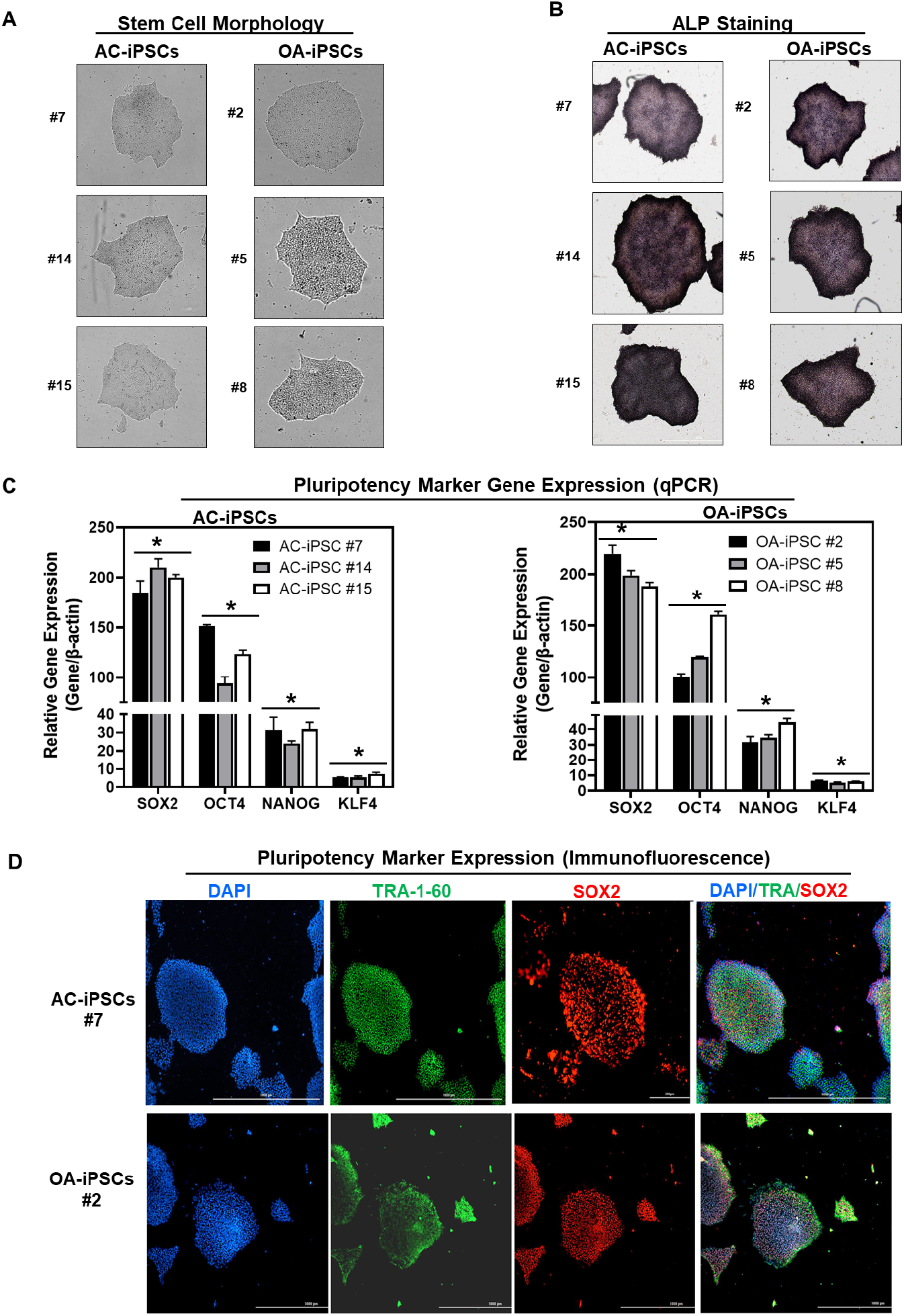
Characterization of iPSCs generated from healthy and OA articular chondrocytes: **(A)** Morphology of the AC-iPSC (#7, #14, #15) and OA-iPSC (#2, #5, #8) colonies in monolayer culture on a 0.1% Geltrex coated plate; **(B)** Alkaline phosphatase (ALP) staining of iPSC colonies showing undifferentiated pluripotent stage; (**C**) Pluripotency for iPSC colonies showing expression of stemness genes. RT-qPCR analyses showed induced expression of canonical stemness genes *SOX2, OCT4, NANOG* and *KLF4* in AC-iPSC and OA-iPSC colonies. β-actin served as the housekeeping gene and internal control. Represented gene expression data is relative to MSCs derived from respective iPSC cells. \**P*≤0.01, as compared to their respective MSCs. (**D**) Immunofluorescence staining of pluripotency markers in AC-iPSCs (#7) and OA-iPSCs (#2) showed expression of surface TRA-1-60 and SSEA-4 antigens in these colonies. DAPI is used as nuclear counterstain showing blue nuclei. Scale bar, 100 μm The online version of this article includes the following source data for figure 1: Figure 1-source data 1. Depicting original raw data related to Figure 1.

### MSCs differentiated from AC-iPSCs and OA-iPSCs exhibit comparable phenotypic features *in vitro*

Differentiation of human iPSCs into mature chondrocytes requires derivation into an intermediate stage termed as mesenchymal progenitors.^13;16;17;22^ Therefore, we generated mesenchymal progenitor intermediate from all three clones of both AC- and OA-iPSCs using our established direct plating method in presence of serum and human recombinant bFGF.^17;22^ MSCs derived from both AC-iPSCs (termed as AC-iMSCs) and OA-iPSCs (termed as OA-iMSCs) displayed similar phenotypic characteristics of spindle-shaped and elongated morphology (**Fig. 2A**). We next performed detailed characterization of iMSCs from both sources to determine their mesenchymal properties. Profiling by qPCR showed significant suppression of stemness genes including *SOX2* and *OCT4,* in both AC-iMSCs and OA-iMSCs as compared to the parental undifferentiated AC-iPSCs and OA-iPSCs respectively (**Fig. 2B**). We also analyzed the expression of marker genes associated with the mesenchymal lineage and our results showed that mRNA expression of *TWIST1* (an epithelial to mesenchymal transition related gene), *COL1A1* (an ECM molecule synthesized by MSCs) and *RUNX1* (a transcription factor expressed in mesenchymal progenitors) was significantly higher in both iMSCs as compared to the pluripotent parental iPSCs (**Fig. 2B**).

**Figure 2:**
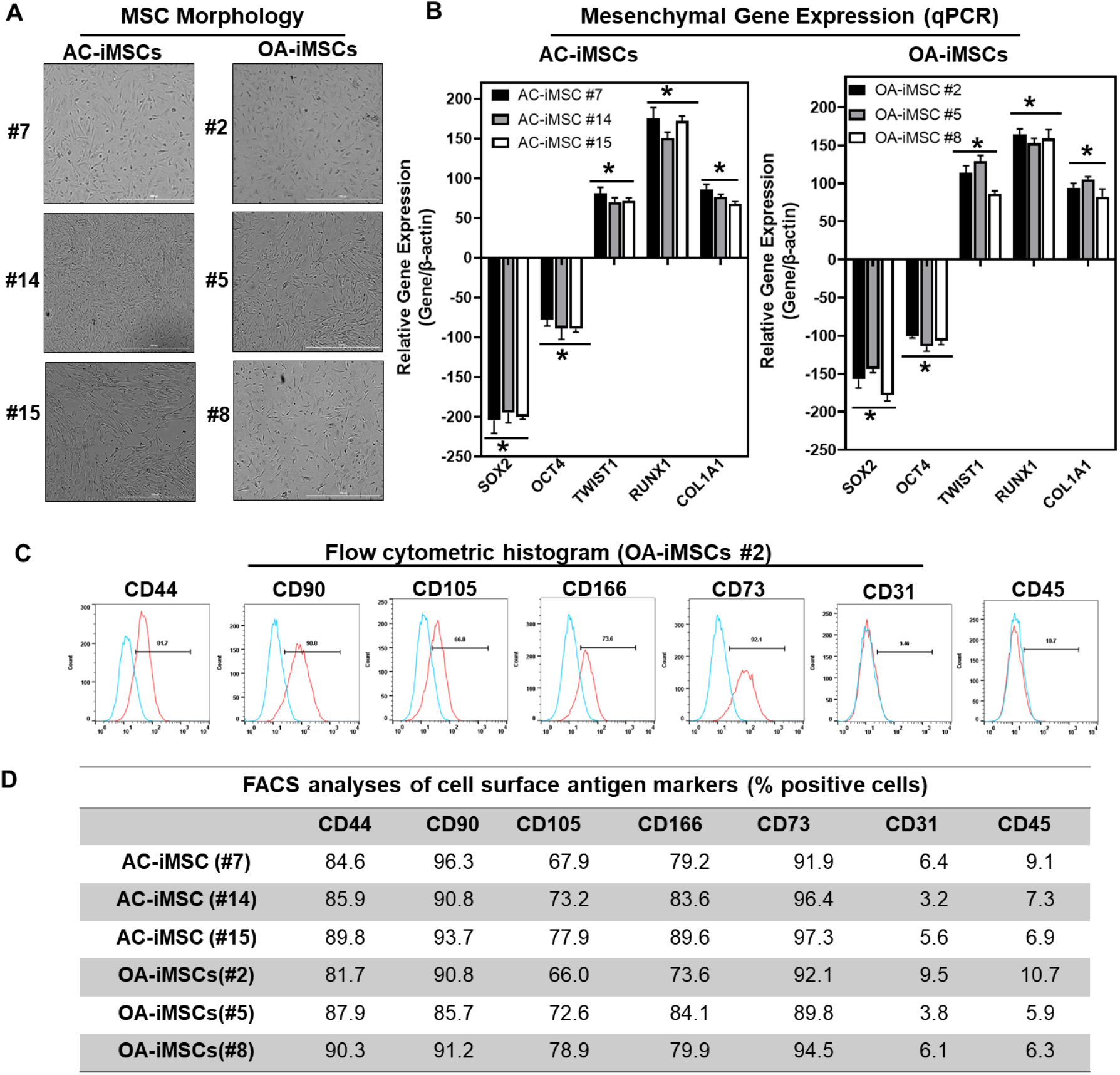
Derivation of iPSC-MSCs (iMSCs) like cells from AC- and OA-iPSCs and characterization of their mesenchymal feature: (A) The morphology of the iMSC-like cells (iPSC–MSC) derived from AC- and OA-iPSC showing elongated spindle shaped cells. Representative images are shown for iMSCs at passage 5-8. Scale bar, 100 μm. (B) Gene expression analyses by qPCR showing significant suppression of pluripotent markers OCT4, and SOX2 and induction of mesenchymal genes TWIST1, COL1A1, and RUNX1 in the AC- and OA-iMSCs relative to their parental iPSCs. β-actin served as the housekeeping gene and internal control. Expression data is represented as fold change relative to respective parental iPSCs. *P≤0.01, as compared to their respective iPSCs. (C) Expression of surface antigens in AC- and OA-iMSCs by flow analysis. Representative flow cytometric histogram showing OA-iMSCs (#2) express markers associated with the mesenchymal phenotype (positive for CD44, CD73, CD90, CD105, and CD166; negative for CD31, and CD45). Red histogram shows antibody-stained population; Blue profile shows negative isotype-stained population. (D) Comparative flow cytometry analyses of AC-iMSCs (#7, #14, #15) and the OA-iMSCs (#2, #5, #8) showing similar cell surface expression profiles. The online version of this article includes the following source data for figure 2: Figure 2-source data 1. Depicting original raw data related to Figure 2.

Consistent with the standard criteria defined by the International Society of Cell and Gene Therapy (ISCT)^18^, immunophenotypic analyses revealed the exhibition of all typical MSC markers in both iMSC progenitors with high expression levels of CD44, CD73, CD90, CD105, and CD166 surface markers (**Fig. 2C**). Conversely, both iMSCs largely lacked expression of the definitive hematopoietic lineage marker CD45, and the endothelial marker CD31. Comparative analysis of these markers in AC-iMSCs and OA-iMSCs showed comparable expression levels suggesting an identical immunophenotype of both iMSCs (**Fig. 2D**). To determine the multipotential of these iMSCs, we performed their trilineage differentiation using *in vitro* adipogenic, osteogenic, and chondrogenic differentiation assays (**Supplementary Fig. 1A, B**). Although both MSCs could clearly form osteoblasts, adipocytes, and chondrocytes, AC-iMSCs displayed enhanced chondrogenic potential as evidenced by increased deposition of Alcian blue positive extracellular matrix compared to OA-iMSCs (**Supplementary Fig. 1B**). Altogether, the data suggest that iMSCs derived from AC-iPSCs and OA-iPSCs exhibit similarities in morphology, immunophenotype, and multipotency as evidenced by *in vitro* differentiation assays for adipocytes and osteoblasts. However, AC-iMSCs displayed increased chondrogenic differentiation as compared to OA-iMSCs.

### AC-iMSCs exhibit enhanced chondrogenic potential *in vitro*

We next evaluated whether AC-iMSCs exhibit higher propensity for chondrogenic differentiation as compared to OA-iMSCs. Chondrogenic differentiation of these iMSCs were examined using our well-established pellet culture method using chondrogenic media in the presence of human recombinant BMP-2 (**Fig. 3A**).^13;16;17;22^ Quantitative PCR analyses of key chondrogenic genes was used to evaluate the potential of AC-iMSCs and OA-iMSCs to produce chondrocytes at days 7, 14, and 21. When compared to the undifferentiated MSC culture (Day 0), induction of *SOX9, COL2A1,* and *ACAN* transcript was significantly increased at day 7, and to a greater extent at day 14 (**Fig. 3C**). Interestingly, mRNA expression of *SOX9, COL2A1,* and *ACAN* were significantly higher in AC-iMSCs as compared to OA-iMSCs at all time points analyzed (Day 7, 14, 21) suggesting that iPSC derived from healthy chondrocytes have a significantly higher chondrogenic potential as compared to OA-iPSC (**Fig. 3C**).

**Figure 3:**
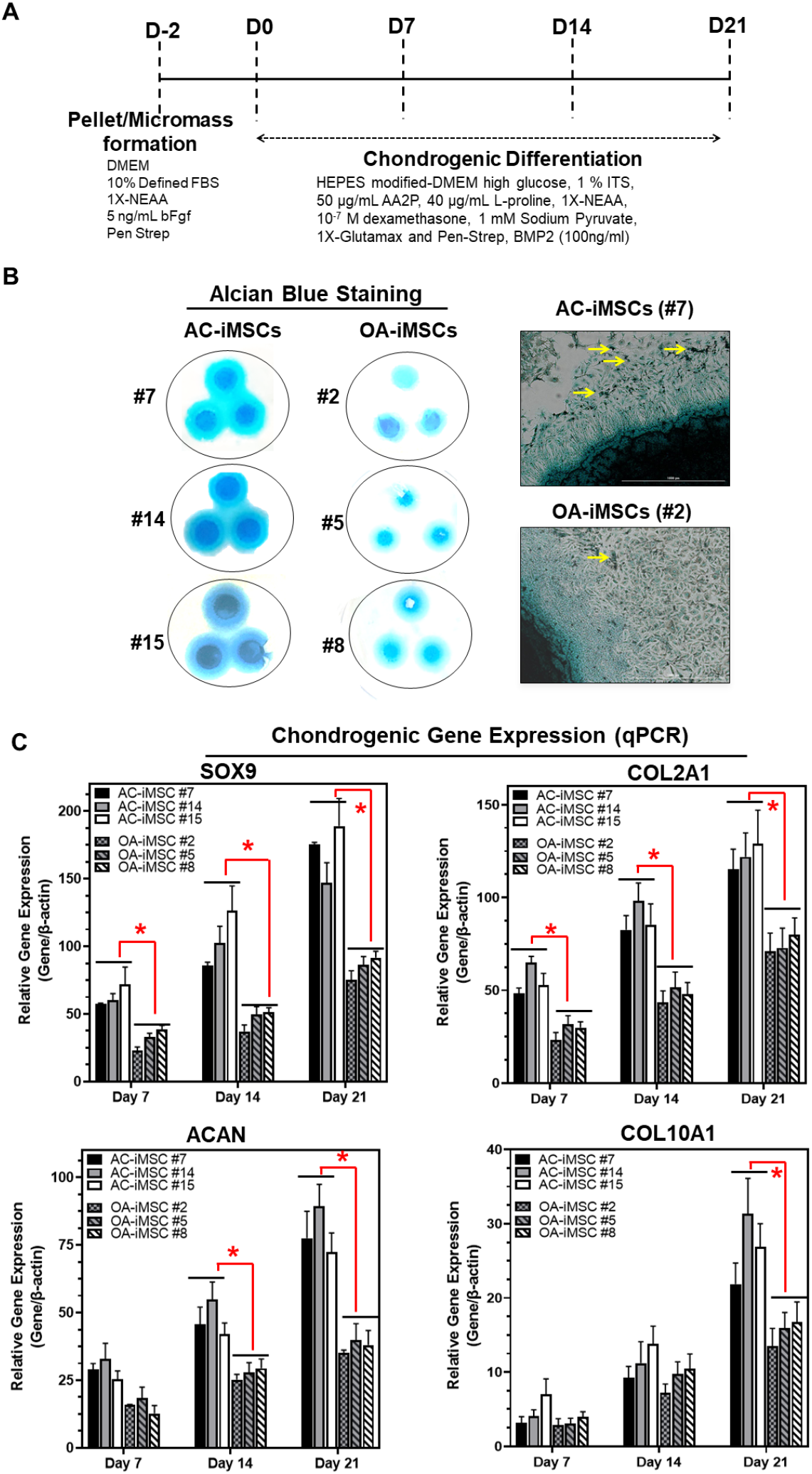
AC-iMSCs exhibit superior chondrogenic potential *in vitro:* **(A)** Schematic showing treatment conditions for *in vitro* chondrogenic differentiation of AC- and OA-iMSCs using pellet and micromass method. (**B**) Chondrocyte differentiation was shown by Alcian blue staining of micromass cultures in serum free chondrogenic media for 21 days with 100 ng/ml BMP2. Alcian blue staining revealed accumulation of sulfated proteoglycans indicating enhanced secretion of matrix in AC-iMSC as compared to OA-iMSCs micromass culture. High magnification images further demonstrated enhanced cellular compaction (yellow arrow) in AC-iMSCs micromass indicating the development of cartilaginous nodules. Scale bar, 100 μm. **(C)** Quantitative PCR analyses of the relative transcript levels of chondrogenic genes *SOX9, COL2A1, ACAN* and hypertrophic gene *COL10A1* in Day 7, 14, and 21 pellet culture of all three clones of AC- and OA-iMSCs. β-actin served as the housekeeping gene and internal control. Values represent fold induction (Mean±SD) relative to control iMSCs (Day 0) from three replicate. *P≤0.01 indicate values are statistically different in OA-iMSCs as compared to their AC-iMSCs at each timepoint. The online version of this article includes the following source data for figure 3: Figure 3-source data 1. Depicting original raw data related to Figure 3.

We also performed chondrogenic differentiation of these iMSCs using high density adherent micromass culture method. 3D-micromass culture of pluripotent stem cells resemble the formation of prechondrogenic mesenchymal condensations and their differentiation into the chondrogenic lineage.^17;32^ Alcian blue staining of Day 21 micromass culture of AC-iMSCs showed densely stained central core surrounded by a diffusely stained outer cellular layer showing increased accumulation of glycoprotein-rich matrix as compared to OA-iMSCs (**Fig. 3B**). Additionally, Alcian blue staining in AC-iMSCs further showed increased cellular outgrowths and cartilaginous nodules, confirming enhanced chondrogenic potential of AC-iMSCs as compared to OA-iMSCs (**Fig. 3B**). These Alcian blue staining showing ECM synthesis are in line with the expression data for the matrix genes. These data further indicate that iMSCs derived from OA chondrocytes showed reduced ECM generation upon chondrogenic differentiation which may be a retention of OA phenotype of original cell source.

### AC-iMSCs exhibit distinct transcriptomic signature during chondrogenic differentiation

To examine the underlying transcriptional programs associated with enhanced chondrogenic potential of AC-iMSCs as compared to OA-iMSCs, we performed RNA-seq analysis. We identified gene expression changes at pan-genome levels in day 21 differentiated chondrocytes from AC-iMSCs (#7) and OA-iMSCs (#5). The volcano plot showed that global gene expression profiles of the chondrocytes at day 21 chondrogenic culture of AC-iMSCs were significantly different from the OA-iMSCs (**Fig. 4A**). This analysis identified 146 genes that were upregulated, and 263 genes that were downregulated in chondrocytes derived from AC-iMSCs (termed as AC-iChondrocytes) as compared to OA-iMSCs (termed as OA-iChondrocytes) (**Fig. 4A**). To validate these findings, we performed quantitative gene expression analysis of a subset of DEGs such as *FOXS1, ADAM12, COL1A1, COL3A1, MATN4,* and *MARK1* during chondrogenic differentiation and analysis confirmed differential expression levels in AC- vs OA-iChondrocytes (**Supplementary Fig. 2**).

**Figure 4:**
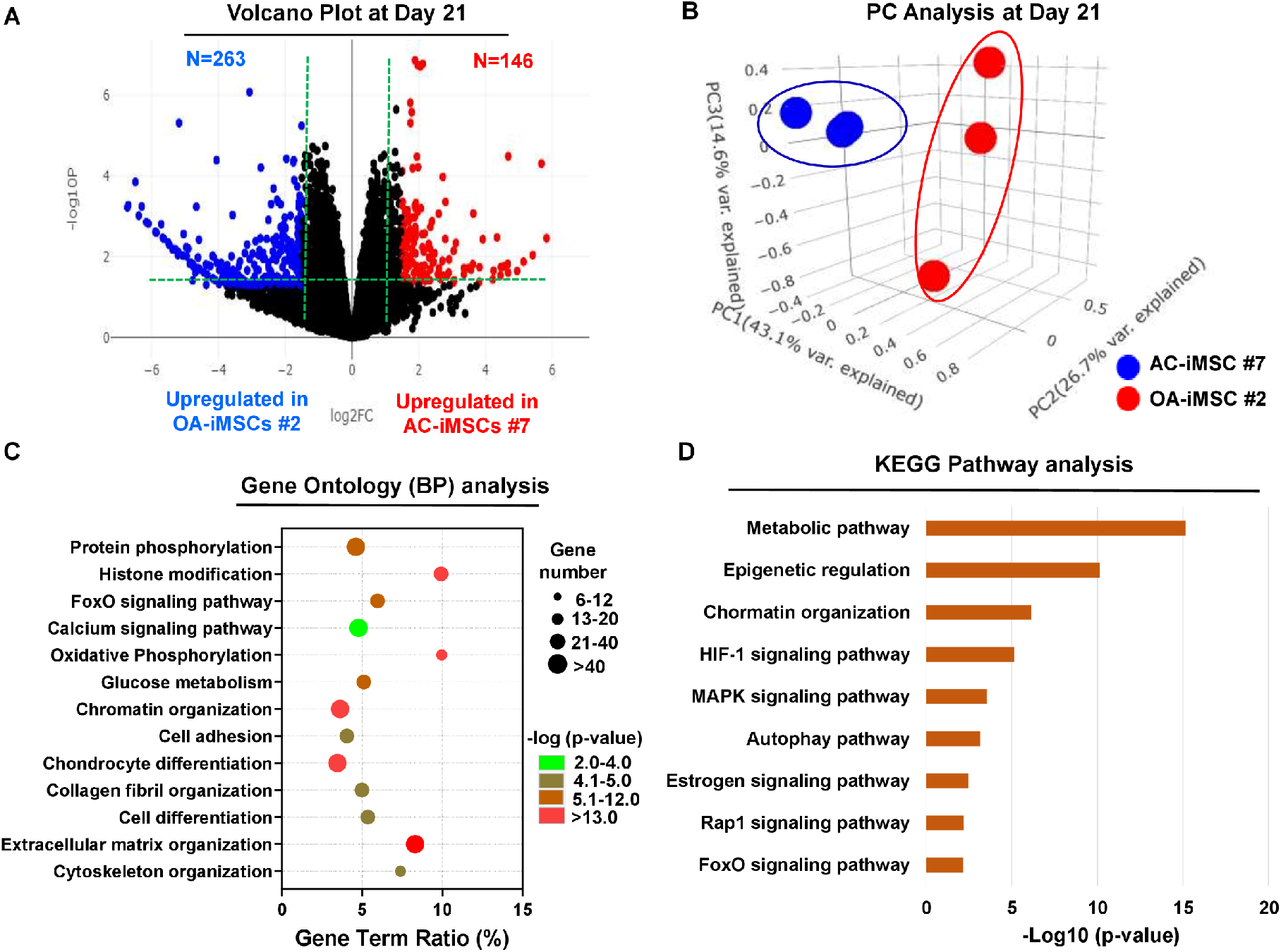
AC-iMSCs during chondrogenic differentiation exhibit distinct transcriptome signature: Bulk RNA-Sequencing was performed during chondrogenic differentiation of AC- vs OA-iMSCs and differential gene expression analyses revealing distinct transcriptomic signature. (**A**) Genes with differential expression levels greater than 2-fold (FDR P value <0.05) were visualized as volcano plot showing differential expression of 406 genes. **(B)** Principal component analysis (PCA) using differentially expressed genes (DEGs) showing segregation of AC-iChondrocytes vs OA-iChondrocytes generated during day 21 chondrogenic differentiation using pellet culture. **(C)** Functional annotation clustering using GO analysis for biological process (BP) using DEGs at day 21 chondrogenic differentiation of AC- vs OA-iMSCs. Y-axis label represents pathway, and X-axis label represents gene term ratio (gene term ratio = gene numbers annotated in this pathway term/all gene numbers annotated in this pathway term). Size of the bubble represents the number of genes enriched in the GO terms, and color showed the FDR P value of GO terms. **(D)** KEGG pathway analysis showing enrichment of molecular pathways contributing to differential chondrogenic potential of AC- vs OA-iMSCs. The online version of this article includes the following source data for figure 4: Figure 4-source data 1. Depicting original raw data related to Figure 4. Figure 4-source data 2. Depicting original raw data related to Figure 4.

Additionally, principal component analyses placed AC- and OA-iChondrocytes in two distinct clusters suggesting that chondrocytes derived from AC-iMSCs were genomically distinct from OA-iMSC derived chondrocytes (**Fig. 4B**).

We next performed functional annotation analyses of these differentially regulated genes to determine the enrichment of GO terms and molecular pathways. Our GO analyses demonstrated significant enrichment of several biological processes in AC-iChondrocytes including histone modification, chromatin organization, oxidative phosphorylation, glucose metabolism, chondrocyte differentiation and ECM organization (**Fig. 4C**). These results suggest that pathways related to ‘Energy Metabolism’ and ‘Epigenetic Regulation’ play an important role in chondrogenic differentiation of AC-iMSCs. We also performed KEGG pathway analysis and results showed that ‘Metabolic Pathways’, ‘Epigenetic Regulation’ and ‘Chromatin Organization’ are the most enriched pathways in AC-iMSCs (**Fig. 4D**). These data suggest that a large proportion of DEGs between AC- and OA-iChondrocytes were involved in ‘Energy Metabolic pathways’ such as oxidative phosphorylation, glucose metabolism and protein phosphorylation and ‘Epigenetic Regulatory pathways’ such as chromatin organization and histone modification. The regulatory genes involved in these pathways such as *HDAC10/11, PRMT6, PRR14, ATF2, SS18L1, JDP2, RUVBL1/2, OGDHL, ALDH2, GCLC, GOT1, HIF1A, COX5A,* and *TRAF6* etc. may create a distinct metabolic and chromatin state in AC-iMSCs which favors its enhanced chondrogenic differentiation.

### AC-iMSCs revealed enrichment of interaction networks related to energy metabolism and epigenetic regulation during chondrogenic differentiation

To determine the functional relationships among genes that were differentially regulated during chondrogenic differentiation of AC- and OA-iMSCs, we performed interaction network analyses. Our analysis identified two major subnetworks distributed in two distinct clusters belonging to energy metabolism and epigenetic regulation suggesting a role for these pathways in chondrogenic differentiation of AC-iMSCs (**Fig. 5A, B**). The ClueGO analysis in metabolic gene network cluster showed enrichment of several energy metabolic pathways such as Glycolysis, Amino acids synthesis, Autophagy and Biosynthesis and anabolic pathways suggesting that multiple metabolic signaling networks in energy metabolism may contribute to enhanced chondrogenic potential of AC-iMSCs (**Fig. 5A**). Similarly, Epigenetic regulator gene network cluster comprise of several pathways related to histone modification, chromatin regulation, histone acetylation and chromatin assembly/disassembly. These data suggest that during chondrogenic differentiation of AC-iMSCs, several chromatin modifiers were activated which may regulate key genes involved chondrogenic differentiation (**Fig. 5B)**.

**Figure 5:**
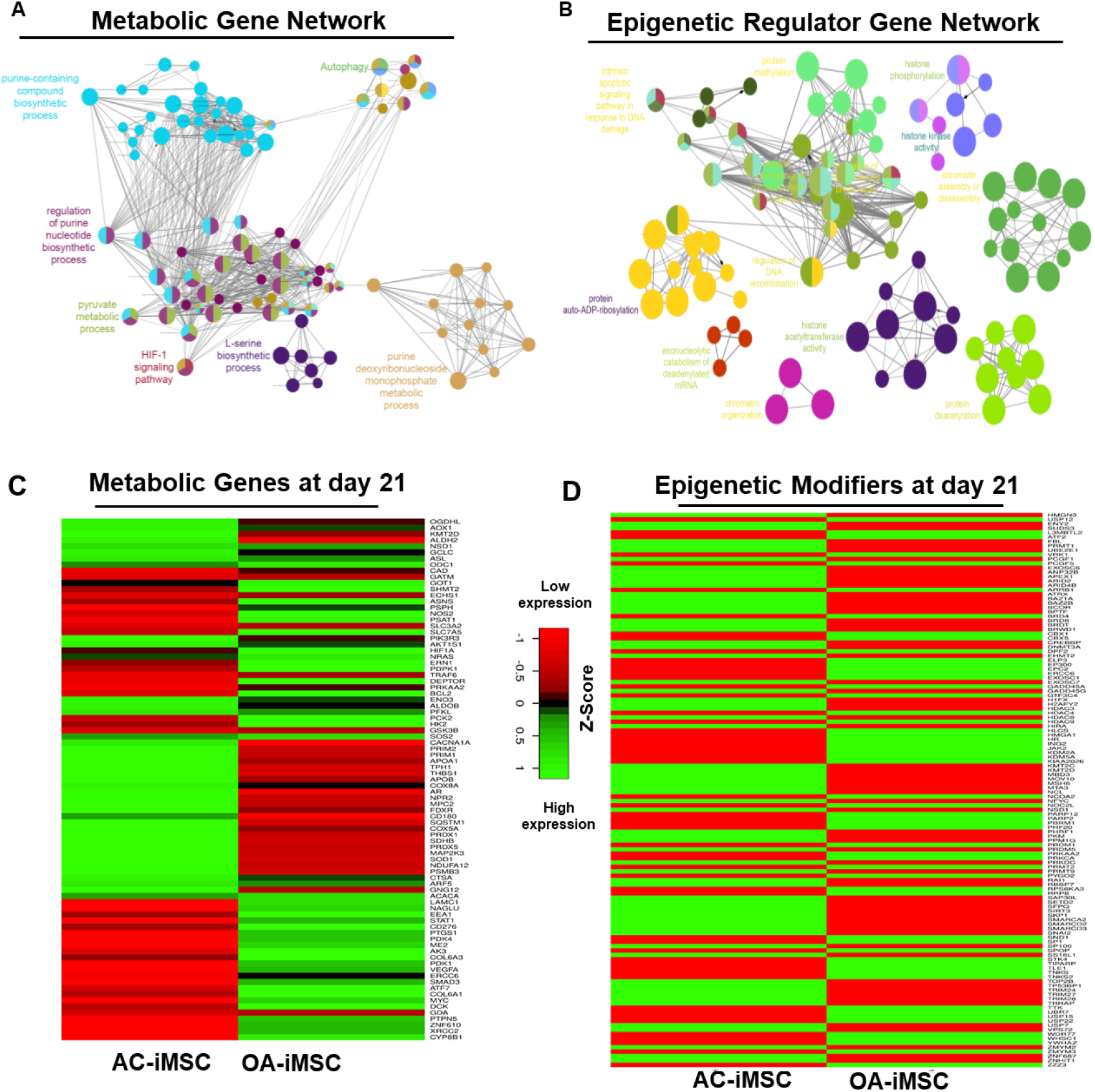
Enrichment of metabolic and epigenetic regulators interaction networks during chondrogenic differentiation of AC-iMSCs: (**A, B**) Interaction network analysis using DEGs at day 21 chondrogenic differentiation of AC- and OA-iMSCs. PPI network of differentially expressed genes in AC-iChondrocytes was constructed using STRING database and visualized by Cytoscape. Pathway enrichment analysis in the interaction network was performed using ClueGO analysis which showed enrichment of pathways related to (**A**) metabolic genes and (**B**) Epigenetic modifiers. Multiple nodes of metabolic and epigenetic regulators were enriched in these interaction networks suggesting the role of these pathways in differential chondrogenic potential. (**C, D**) Differential expression analyses of the genes involved in these enriched pathways related to energy metabolism and epigenetic regulators. The gene expression was visualized using Heatmap analysis for differentially expressed genes related to (**C**) energy metabolism and (**D**) epigenetic regulators. Expression values for each gene (row) were normalized across all samples (columns) by Z-score. Color key indicates the intensity associated with normalized expression values. Green shades indicate higher expression and red shades indicate lower expression. The online version of this article includes the following source data for figure 5: Figure 5-source data 1. Depicting original raw data related to Figure 5. Figure 5-source data 2. Depicting original raw data related to Figure 5. Figure 5-source data 3. Depicting original raw data related to Figure 5.

To further implicate the role of ‘energy metabolism’ and ‘epigenetic regulator pathways’ in differential chondrogenic potential, we analyzed the expression profile of genes involved in these pathways during chondrogenic differentiation of AC- and OA-iMSCs. The heatmap analysis during terminal chondrogenic differentiation (Day 21) showed that the expression profile of various metabolic and epigenetic regulator genes exhibits differential expression in AC- vs OA-iChondrocytes. (**Fig. 5C, D**). Moreover, the expression profile for metabolic and epigenetic factor genes correlates well with chondrogenic differentiation of these iMSCs further suggesting the importance of these pathways in enhanced chondrogenic potential of AC-iMSCs. Altogether, these data suggest that metabolic and epigenetic regulatory pathways play a role in chondrogenic potential of AC-iMSCs.

### AC-iMSCs at the undifferentiated state showed distinct expression of genes involved in energy metabolism and epigenetic regulation

The data in **Figure 5C, D** show that during chondrogenic differentiation, AC-iMSCs exhibit differential expression for the metabolic and chromatin regulator genes. We next examined whether this differential gene expression profile was intrinsic to AC-iMSCs or acquired during the process of chondrogenic differentiation. To this end, we performed transcriptomic analyses at various stages of chondrogenic differentiation of AC- and OA-iMSCs. Volcano plot analysis identified that AC- and OA-iMSCs at undifferentiated steady-state (Day 0) exhibited differential expression at pan-genome level with >800 differentially expressed genes (**Fig. 6A**). We next focused our analysis on the expression of genes involved in energy metabolic and epigenetic regulator pathways. Similar to the level observed at terminal differentiation stage, our analysis revealed that metabolic and chromatin regulator genes also showed significant differences between both cell types at the uncommitted mesenchymal state (**Fig. 6B, C**). When compared to OA-iMSCs, the AC-iMSCs expressed higher levels of several metabolic gene involved in glycolysis, amino acid synthesis, autophagy, and anabolic pathways such as *ALDOB, CD180, SQSTM1, ENO3, AOX1, KMT2D, COX5A, PRDX1, SDHB, ALDH2* etc (**Fig. 6B**). Moreover, differential expression of multiple chromatin modifiers including histone modifiers (eraser, reader and writers) and chromatin remodeling factors such as *JDP2, RUVBL1/2, MYBBP1A, HDAC10, HDAC11, USP12, L3MBTL2, MUM1* etc was also observed (**Fig. 6C**). Further, differential expression patterns of several epigenetic modifiers at the MSC stage (day 0) were retained at the chondrocyte stage (day 21). For example, *ARID4B, BRD4, HDAC4, HDAC9, KDM5A, KMT2C* etc showed differential expression between OA- and AC-iMSCs at both day0 and day 21 stage of chondrogenic differentiation. These results suggest that differential expression of genes associated with energy metabolism and epigenetic regulation between healthy and OA conditions first occurs at the MSC stage, prior to their overt differentiation to the chondrogenic lineage. Thus, differences in the chondrogenic potential of AC-versus OA-iMSCs may be associated with differences in expression of metabolic and chromatin modifier genes which influence the chondrogenic capacity of these MSCs.

**Figure 6:**
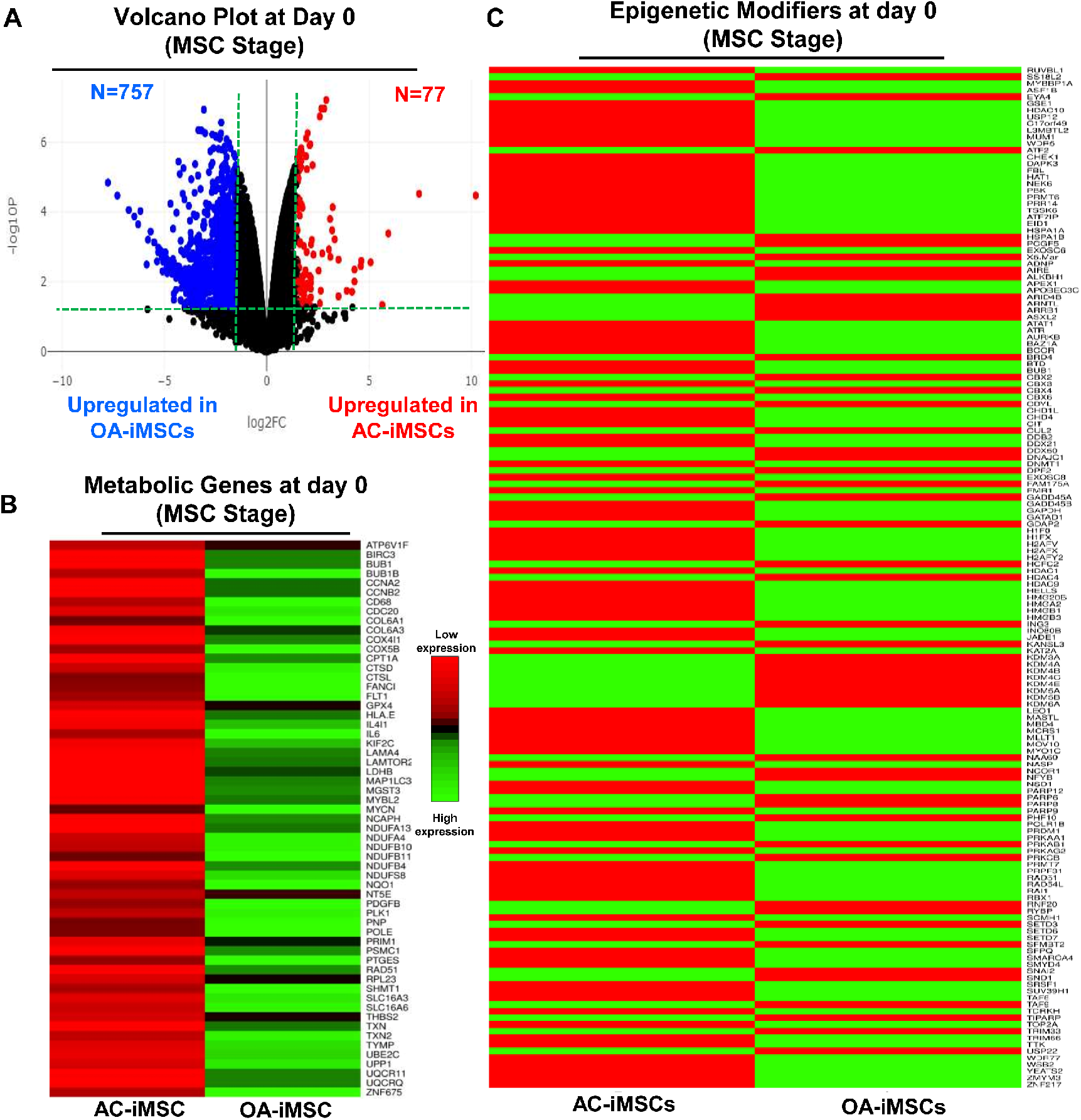
AC-iMSCs at undifferentiated state showed distinct expression of genes involved in metabolic and epigenetic regulators: **(A)** Differential gene expression analyses of AC- and OA-iMSCs at day 0 (start of chondrogenic differentiation) showing distinct transcriptomic signature. Genes with differential expression levels greater than 2-fold (FDR P value < 0.05) were visualized as volcano plot showing differential expression of 834 genes. (**B, C**) Pathway analysis was performed in 834 DEGs to show the enrichment of pathways related to metabolism and Epigenetic modifiers. Heatmap was used to show expression of the genes related to (**B**) energy metabolism and (**C**) epigenetic regulators. Expression values for each gene (row) were normalized across all samples (columns) by Z-score. Color key indicates the intensity associated with normalized expression values. Green shades indicate higher expression and red shades indicate lower expression. The online version of this article includes the following source data for figure 6: Figure 6-source data 1. Depicting original raw data related to Figure 6. Figure 6-source data 2. Depicting original raw data related to Figure 6.

### Genetically distinct characteristic of AC- and OA-iMSCs were imprint of original cell sources from healthy and OA-chondrocytes

Our data as above (**Fig. 2A-E** and **Fig. 6A-C**) indicated that although AC- and OA-iMSCs exhibit similar morphologic and immunophenotypic characteristics, they are genetically distinct populations that displayed varying efficiencies for chondrogenic differentiation. We therefore postulated that differences in the metabolic and chromatin modifier gene expression patterns observed in OA-iMSCs as compared to AC-iMSCs are attributed to their initial disease status. To explore this, we analyzed the expression profiles of metabolic and chromatin modifier genes from multiple sources of healthy and OA-chondrocytes. Thus, we analyzed publicly available RNA-seq data performed on healthy and OA cartilage tissues (GSE114007).^28^ This analysis revealed that the expression profiles of a large number of metabolic and chromatin modifier genes are differentially expressed in human AC-versus OA-cartilages (**Fig. 7A, B**). Similarly, we detected differential expressions of key metabolic and epigenetic modifiers in our unbiased datasets from AC-versus OA-iMSCs (uncommitted stage, day 0), suggestive of the persistence of a cellular memory of disease even after reprogramming. Several chromatin modifiers such as *HAT1, HDAC10, HDAC11, PRMT6, JDP2, ATF2, ATF7, WDR5* etc. which showed differentiation expression in healthy and OA cartilage also showed retention of differential pattern in OA- vs AC-iMSCs. Together our data suggest that a retained memory of disease during stem cell reprogramming affected the chondrogenic differentiation potential of OA-iMSCs.

**Figure 7:**
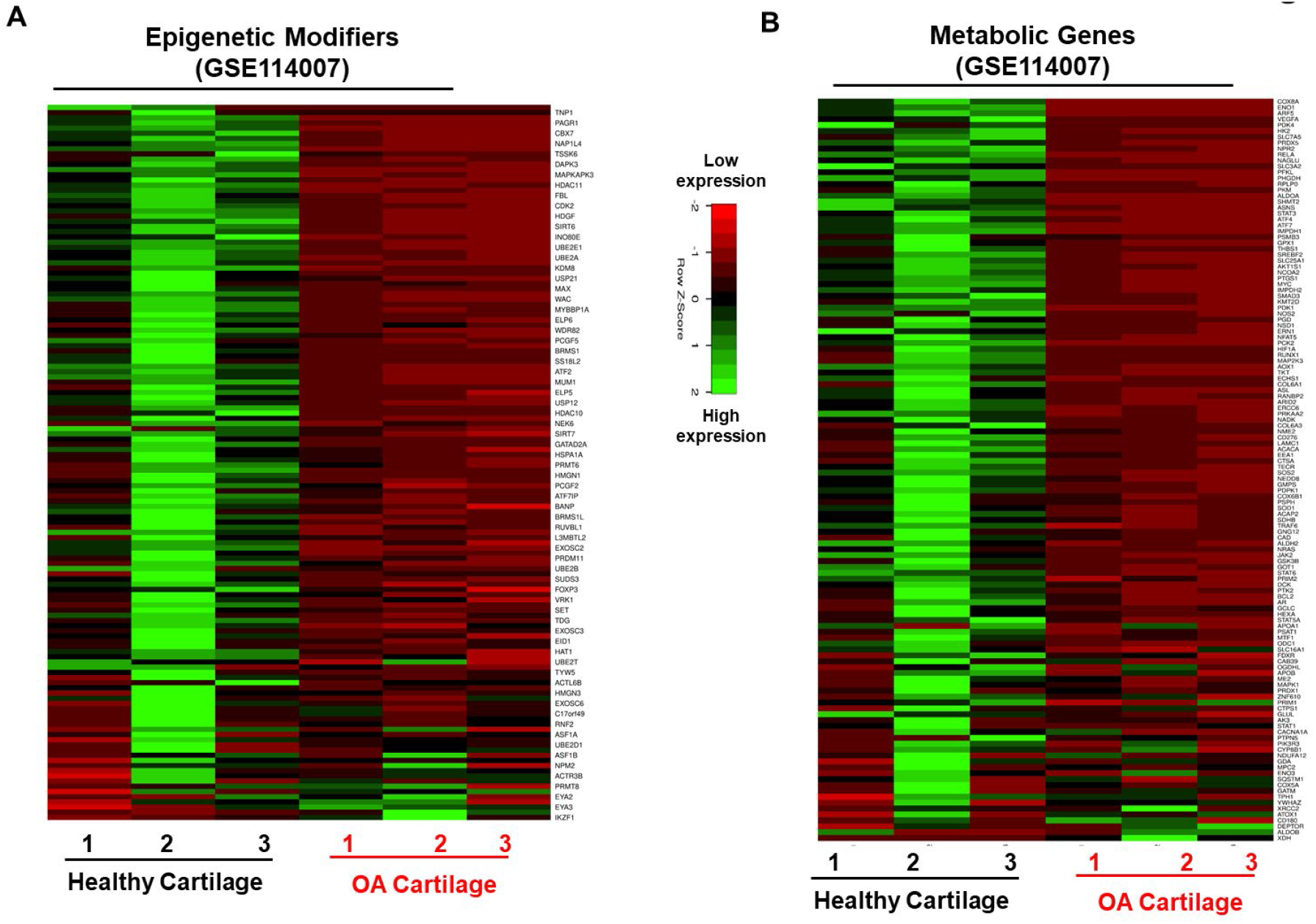
Genetically distinct characteristic of AC- and OA-iMSCs were imprint of original cell sources from healthy and OA-chondrocytes: The expression of genes involved in energy metabolism and epigenetic modifiers was performed by analysis of publicly available RNA-seq data performed on healthy and OA cartilage tissues (GSE114007). Gene expression was visualized by heatmap analysis for (**A**) epigenetic modifiers and (**B**) metabolic genes. Expression values for each gene (row) were normalized across all samples (columns) by Z-score. Color key indicates the intensity associated with normalized expression values. Green shades indicate higher expression and red shades indicate lower expression. The online version of this article includes the following source data for figure 7: Figure 7-source data 1. Depicting original raw data related to Figure 7. Figure 7-source data 2. Depicting original raw data related to Figure 7.

## DISCUSSION

iPSCs are viewed as promising cell-based therapeutics for the repair of tissues lacking intrinsic regenerative capacity, including articular cartilage. Multiple studies have cautioned that safe and effective application of iPSCs based therapeutics will require careful consideration of the cellular origins of iPSCs.^33 34^ Although reprogramming of somatic cells to iPSCs involves extensive modification of the epigenetic landscape, the reprogrammed cells can retain an epigenetic memory of the cell type of origin, thus affecting lineage differentiation propensity.^5;10;35^ In addition to donor cell type, key questions over the influence of the health status of the parental somatic cells used for reprogramming remain unresolved. Thus, the present study was designed to determine whether the health status of donor human articular chondrocytes influences the regenerative potential of the derived iPSCs. Using iPSCs generated from healthy and OA chondrocytes, we report that reprogramming efficiency to pluripotency was largely equivalent between the two sources. However, OA-iPSCs showed a significantly reduced capacity for chondrogenic differentiation as compared to AC-iPSCs, indicating that the pathogenic condition of the donor chondrocytes negatively affected the chondrogenic differentiation potential of OA-iPSCs. Our data suggest that reprogramming does not reset the health status of OA articular chondrocytes, but rather supports the existence of a memory of disease in iPSCs derived from OA cartilage.

A plethora of studies over the last 15 years have determined that cells from almost any tissue can be used to generate human iPSCs, which can then be differentiated to a variety of specialized cells. However, human iPSCs generated from disparate cell types have not displayed equivalent capacities for differentiation to specialized cell types.^10^ Seminal studies using iPSCs derived from myeloid cells, hematopoietic cells, and insulin-producing β-cells revealed a biased lineage differentiation attributed to residual DNA methylation signatures that influence cell fate commitment. For example, when compared to isogenic non-beta cell-derived iPSCs, beta cell-derived iPSCs maintained an open chromatin structure at key beta-cell genes, leading to an increased capacity to efficiently differentiate into insulin-producing cells.^4^ Thus, iPSCs appear to have an epigenetic memory for the tissue of origin. We have previously generated iPSCs from multiple cell types including human skin fibroblasts, umbilical cord blood, and normal healthy chondrocytes using the same reprogramming strategy.^13^ Using multiple chondrogenic differentiation assays, our earlier findings demonstrated that iPSC derived from chondrocytes showed enhanced matrix formation and chondrogenic gene expression, suggesting that the tissue of origin also impacted the chondrogenic potential of human iPSCs.^13^ Similarly, a previous report also demonstrated the differential chondrogenic capabilities of iPSCs derived from dermal fibroblasts, peripheral blood mononuclear cells, cord blood mononuclear cells and OA fibroblast-like synoviocytes.^8^

Although it is now well-documented that the tissue of origin can affect the differentiation potential of iPSCs^10^, it is not known whether the health status of same tissue affects the regenerative potential of its derived iPSCs. A combination of genetic and non-genetic factors, including advanced age, mechanical trauma and inappropriate joint loading, and inflammation contribute to the development of OA.^36–38^ It is well established that human OA articular chondrocytes exhibit phenotypic, functional and metabolic changes, as well as altered epigenetic patterns.^39^ Thus, we speculated that a retained epigenetic memory of iPSCs is not only specific to the tissue of origin but also to the diseased status. Using pluripotency as a reliable tool, our novel data demonstrated significant differences in the chondrogenic capability of AC- vs OA-iPSCs. Our data indicate that OA cartilage derived iPSCs retained functional and molecular characteristics of OA pathogenesis. Chondrogenic pellets generated using OA-iPSCs showed relatively smaller size and reduced chondrogenic gene expression as compared to that from healthy iPSCs (AC-iPSCs) Expression of the trio of SOX genes *(SOX9, SOX5* and *SOX6)* was significantly lower in OA-iPSC-derived chondrogenic pellets. The expression of chondrogenic genes under control of SOX9, such as *COL2A1* and *ACAN* were also lower than that AC-iPSC-derived pellets. Since expression of *COL2A1* and *ACAN* are usually lower in OA cartilage, the finding of reduced chondrogenic genes in OA-iPSCs during chondrogenesis suggest the imprints of disease pathogenesis in OA-iPSCs. Similarly other cartilage matrix genes such as *COMP, MATN4, PRG4* and *COL11A2* were lower in OA-iPSC derived chondrocytes. Interestingly, expression of these genes was also reported to be decreased in OA^40^ further suggesting the recapitulation of memory of disease in OA-iPSCs as compared to AC-iPSCs. Based on these findings, human stem cell models of OA using iPSCs may provide the unique opportunity to model OA disease changes, to uncover mechanisms of disease development, and to identify molecular targets for therapeutic intervention.

We next identified how a memory of cartilage pathology in OA is transmitted from the original somatic cells to iMSCs and finally to the chondrocytes. A nonbiased, high-throughput RNA-sequencing approach was used to define the pan transcriptome changes during iPSC stage-specific differentiation. Our global transcriptome data showed skewed expression of epigenetic regulators, and metabolism-associated molecular pathways in AC- vs OA-iMSCs, suggesting a transcriptional memory of disease mechanisms in OA-iPSCs. Recent studies showed cellular metabolism as a key driver of cell-fate changes which has intrinsic links with epigenetic modifications of chromatin during development, disease progression, and cellular reprograming.^41^ Our data suggests that AC- and OA-iMSCs differ in the expression of a plethora of metabolic genes which finally influence the cells metabolism and thus chondrogenic differentiation. While cell metabolism is closely linked to chondrogenic differentiation, in-depth metabolomic studies are needed to determine how metabolic heterogeneity of AC- and OA-iPSCs impact chondrogenic differentiation and regenerative potential of cartilage tissue. While recent discovery^41^ demonstrated the interaction between energy metabolite and epigenetic modifiers, a detailed future investigation warrants to determine how cellular metabolism wired the epigenetic modification and influence the cellular transitions associated with cartilage development.

Although an apparent memory of disease can impact the chondrogenic capabilities of OA-iPSCs, we did not detect differences in stemness genes between AC and OA lines. These data indicate that transcriptome level differences were notable only upon initial differentiation towards uncommitted mesenchymal progenitors (iMSC stage). We do not know whether the functional and molecular alteration in OA-iMSCs represent a transient or stable phenomenon. Functional studies, coupled with comprehensive analyses of epigenetic landscapes will be necessary to address whether the observed memory of disease (epigenetic and metabolic) is a stable imprint of the original cellular phenotypes, or could be erased by serial reprogramming. Moreover, does preservation of an epigenetic memory of cartilage disease in iPSCs occur at the DNA methylation level, and if so, what are the OA-associated loci? Further, it is not clear whether memory of disease is a phenomenon observed only at early passages after pluripotency induction or can be attenuated by continuous passaging.

In the present study, we addressed for the first time that differential chondrogenic potential of AC- and OA-iPSCs could be attributed to differences in transcriptome level changes in the epigenetic modifiers and energy metabolic genes. The expression profile of several chromatin modifiers belonging to the family of histone readers, writers, and erasers such as *FBL, PRMT1, UBE2E1, VRK1, PCGF1, USP12, HMGN3, HDAC3, HDAC8, BRDT, ARID2,* and *HMGN3* were significantly different between AC- and OA-iMSCs. In addition, several metabolic genes such as *AOX1, OGDHL, GATM, KMT2D, ALDH2, GOT1, SLC3A2* and *ECHS1* also showed differential expression pattern between AC- and OA-iMSCs. Several studies previously showed that metabolic genes and metabolites are involved in the regulation of histone acetylation and chromatin modification indicating that importance of chromatin and metabolites in physiological function of the cells.^42;43^ Future studies using genome editing approaches coupled with metabolomics and chromatin mapping approaches will be required to determine the biological roles of these identified chromatin modifiers and metabolic regulators in chondrogenic differentiation of iMSCs. Further correlation of chondrogenic differentiation potential of iMSCs derived from chondrocytes from multiple donors, and with varying grades of OA severity will further help establish the concept of epigenetic memory of disease and determine the influence epigenetic and metabolic imprints on cartilage repair and regenerative medicine.

## MATERIALS AND METHODS

### Key resources table

| **Reagent type (species) or resource** | **Designation** | **Source or reference** | **Identifiers** | **Additional Information** |
| --- | --- | --- | --- | --- |
| Biological sample (Human) | AC-iPSCs | Generated at corresponding author lab at UConn Health (Guzzo MR et al 2012) |  |  |
| Biological sample (Human) | OA-iPSC | Generated at corresponding author lab at UConn Health |  |  |
| Biological sample (Human) | AC-iMSCs | Derived from AC-iPSCs |  |  |
| Biological sample (Human) | OA-iPSC | Derived from OA-iPSCs |  |  |
| Antibody | FITC Mouse Anti-Human CD44 | BD-Biosciences | 347943, RRID:AB_400360 | 1:100 for Flow cytometry |
| Antibody | PE Mouse Anti-Human CD73 | BD-Biosciences | 550257, RRID:AB_393561 | 1:100 for Flow cytometry |
| Antibody | FITC Mouse Anti-Human CD90 | BD-Biosciences | 555595, RRID:AB_395969 | 1:100 for Flow cytometry |
| Antibody | PE Mouse Anti-Human CD166 | BD-Biosciences | 559263, RRID:AB_397210 | 1:100 for Flow cytometry |
| Antibody | FITC Mouse anti-Human CD105 | BD-Biosciences | 561443, RRID:AB_10714629 | 1:100 for Flow cytometry |
| Antibody | FITC Mouse Anti-Human CD31 | BD-Biosciences | 555445, RRID:AB_395838 | 1:100 for Flow cytometry |
| Antibody | FITC Mouse Anti-Human CD45 | BD-Biosciences | 347463, RRID:AB_400306 | 1:100 for Flow cytometry |
| Antibody | FITC Mouse IgG1, $\kappa$ Isotype Control | BD-Biosciences | 349041, RRID:AB_400397 | 1:100 for Flow cytometry |
| Antibody | PE Mouse IgG1, κ Isotype Control | BD-Biosciences | 555749, RRID:AB_396091 | 1:100 for Flow cytometry |
| Commercial assay or kit | High-capacity cDNA Reverse | Transcription Kit Applied Biosystems | 4368814 |  |
| Commercial assay or kit | Powerup SYBR green Mix | Thermo-Fisher | A25742 |  |
| Chemical compound and drugs | Trizol | Thermo-Fisher | 15596026 |  |
| Chemical compound and drugs | DMEM | Thermo-Fisher | 11965092 |  |
| Chemical compound and drugs | Recombinant Human FGF-basic | Peprotech | 100-18B |  |
| Chemical compound and drugs | Non-Essential Amino Acids Solution | Thermo-Fisher | 11140050 |  |
| Chemical compound and drugs | HyClone™ Fetal Bovine Sera Defined | VWR | 16777-006 |  |
| Chemical compound and drugs | Pen Strep | Thermo-Fisher | 10378016 |  |
| Chemical compound and drug | L-Ascorbic Acid | Sigma | A4544 |  |
| Chemical compound and drug | Glutamax 100 X | Gibco | 35050-061 |  |
| Chemical compound and drug | Dexamethasone | Sigma | D2915 |  |
| Chemical compound and drug | L-Proline | Sigma | P0380 |  |
| Chemical compound and drug | Insulin-Transferrin-Selenium | Thermo-Fisher | 41400045 |  |
| Software and algorithm | Prism | GraphPad | RRID:SCR_002798 |  |
| Software and algorithm | DESeq2 | Bioconductor | DESeq2, RRID:SCR_015687 |  |

### iPS cell induction and culture

We have previously described the generation iPSCs reprogramming from human chondrocytes isolated from normal healthy cartilage (AC-iPSCs).^13^ These iPSCs were fully reprogrammed and detailed characterization of pluripotency were performed previously using various methods including molecular, cytochemical, cytogenic and *in vitro* and *in vivo* functional analyses.^13^ Using similar methods, we derived and characterized iPSCs from OA chondrocytes (OA-iPSCs). The OA-iPSCs were generated at UConn Health with IRB approval. We procured surgical discards from a 77-year-old female patient undergoing knee joint replacement surgery at our clinic. Chondrocytes were harvested from remaining, OA-affected cartilage at the tibia plateau. OA-derived iPSCs were generated using polycistronic STEMCAA lentiviral vector (as described in our previous publication).

We used three clones from each of the AC-iPSCs (clone #7, #14, and # 15) and OA-iPSC (clone #2, #5 and #8) to ensure that our data are not clone specific. The iPSC colonies were maintained in undifferentiated pluripotent state by culturing the cells under feeder free conditions on 0.1% Geltrex® (Peprotech) coated culture plates. For routine expansion, iPSCs colonies were passaged after reaching 70% confluency using treatment of ReLeSR™ reagent (StemCell Technologies) and cultured in new 6-well plate using mTeSR™ plus medium supplemented with 10 μM Y-27,632 Rock inhibitor (StemCell Technologies). Pluripotency of all lines was established by analyzing the expressions of canonical stemness genes *(SOX2, NANOG, OCT4, KLF4)* using qPCR assay as described previously.^14^ Full list of primers is listed in Table 1. We also performed immunofluorescence staining for pluripotency markers in these iPSC colonies using Pluripotent Stem Cell 4-Marker Immunocytochemistry Kit (Thermo Fischer Scientific) as per manufacturer’s instruction and fluorescence were imaged using fluorescence microscopy (BioTek Lionheart LX Automated Microscope) as described previously.^15^ Alkaline phosphatase (ALP) staining was also performed for pluripotency characterization using TRACP & ALP double-stain Kit (Takara) following manufacturer’s instructions. ALP-positive colonies were imaged using Automated Microscope (BioTek Lionheart LX).

**Table 1:**
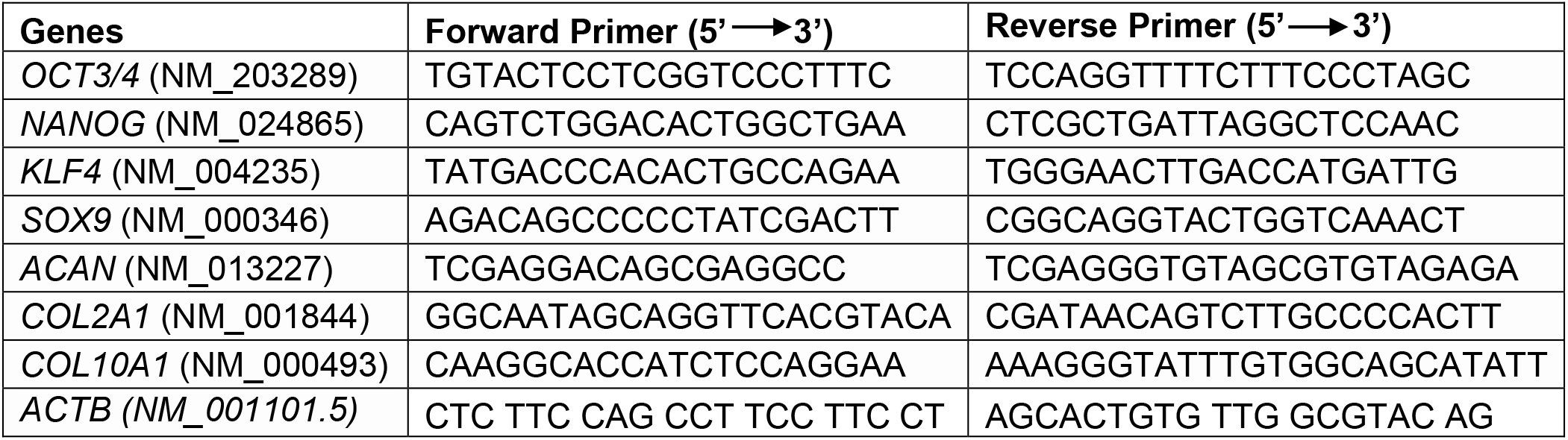
Primer Sequences.

### Derivation of mesenchymal progenitor cells from AC- and OA-iPSCs

Differentiation of iPSCs into chondrocytes requires an intermediate state which we termed as uncommitted mesenchymal progenitor cells or mesenchymal stem like cells (MSCs). The differentiation of iPSCs into MSCs was performed using our established direct plating method as described previously.^16;17^ Briefly, cell suspensions of iPSC colonies (P15-17) were prepared using accutase treatment followed by seeding onto gelatin coated culture plate using MSC growth medium consisting of DMEM-High Glucose (Gibco), 10% defined fetal bovine serum (FBS; Hyclone), 1% nonessential amino acids (NEAA), 1X penicillin-streptomycin, and 5 ng/ml rhbFGF (Peprotech). After 2-3 passage onto non-coated plates, the heterogenous cultures acquired the iPSC-MSC-like homogenous, fibroblast-like morphology which was termed as iPSC derived MSCs (referred as iMSCs). For routine expansion, AC-iPSC, and OA-iPSC derived MSCs (AC-iMSCs and OA-iMSCs) were plated at density of 0.3-0.4×10^6^ cells per 100mm culture dish and maintained in MSC growth media. The characterization of the MSC like feature was performed using gene expression analysis of mesenchymal genes by qPCR assay as described previously.^15^

### Flow characterization of mesenchymal progenitor cells (iMSCs)

Immunophenotyping analysis for cell surface markers was performed as defined by the International Society for Cell & Gene Therapy (ISCT) for the minimal criteria of MSCs.^18^ Surface staining of MSCs markers were performed using labeled anti-human antibody against CD73, CD95, CD105, CD44, CD45, CD31, CD29 using method described previously.^19–21^ Isotype-matched control (IgG1-PE and IgG2b-FITC) were used for identifying nonspecific fluorescence. Cells were acquired using BD FACSAria™ using FACS Diva software (Becton–Dickinson). For each analysis, minimum of 20,000 cells was acquired and data was analyzed using FlowJo Software as described previously.^15^

### Chondrogenic differentiation of iMSCs

We performed chondrogenic differentiation of iMSCs (P18-22) in 3D high density culture conditions using pellet suspension and micromass adherent method using our established protocol as described previously. ^13;16;17;22^ Briefly, for pellet culture, single cell suspension of AC-iMSCs and OA-iMSCs culture was performed using 0.25% Trypsin-EDTA and 0.5×10^6^ cells were placed in 15-ml polypropylene tubes and centrifuged at 300g for 5 minutes to pellet the cells, and finally cultured in MSC growth medium in CO2 incubator at 37°C and 5% CO_2_ for 1 day. Twenty-four hours after pellet formation, the culture media was replaced with chondrogenic media consisting of DMEM-High Glucose media (Gibco), 1% ITS+ premix, 40 μg/ml L-proline, 1mM sodium pyruvate, 1X nonessential amino acids, 1X Glutamax, 50 μg/ml ascorbic acid 2-phosphate, and 0.1μM dexamethasone, 1X-penicillin/streptomycin and human recombinant BMP-2 (100 ng/ml, Peprotech).^16^ Chondrogenic media and growth factor were changed every other day until the end of 21 days of chondrogenic differentiation. Cell pellets were harvested at 0, 7, 14 and 21 days of differentiation and analyzed for gene expression using SYBR™Green qPCR assay.

Chondrogenic differentiation was also performed using adherent micromass method.^16^ The micromass of AC-iMSCs and OA-iMSCs was prepared by culturing the cells in high density (25 ×10^4^ cells per 10μl drop) in 6-well culture plates. Immediately after seeding the micromasses, MSC growth medium was carefully added dropwise from the edges of the plate to prevent dehydration of micromass. These micromasses were incubated for 4-6 hrs at 37°C in 5% CO2, and then supplemented with 2 ml of MSC growth medium and cultured for 24 hrs. Then MSC growth media was replaced with chondrogenic media and differentiation was continued for 21 days. Micromass culture were harvested at different days of chondrogenic differentiation and processed for either RNA isolation or fixed with formalin for Alcian blue staining. Formalin-fixed micromass cultures were stained with 1% Alcian Blue in acetic acid, pH 2.5 and proteoglycans levels were measured by imaging the blue colonies using automated microscope (BioTek Lionheart LX).

### Osteogenic and Adiopogenic differentiation of iMSCs

To establish the multilineage potential of the iMSCs, we next assessed the ability of AC- and OA-iMSCs to differentiate into osteogenic and adipogenic lineages *in vitro.* Osteogenesis of iMSCs was induced by culturing 10,000 cells per well of 24-well plate using osteogenic medium consisting of DMEM supplemented with 1mM sodium pyruvate, 0.1μM dexamethasone, 50 μg/ml ascorbic acid 2-phosphate, 10mM β-glycerophosphate, 10% FBS and 1X penicillin/streptomycin for 21 days. At end of 21 days culture, cells were fixed with formalin and stained for Alizarin Red solution to visualize calcium deposits as described previously.^13;16;17^

Additionally, to induce adipogenesis, the iMSCs were seeded at 10,000 cells per well in 24-well plate and cultured for 21 days in presence of adipogenic media consisting of DMEM-high glucose supplemented with 10% FBS, 1mM sodium pyruvate, 1μM dexamethasone, 10 μg/ml insulin, 0.5mM isobutylmethylxanthine, 200μM indomethacin and 1X penicillin/streptomycin. Adipogenesis was measured by Oil-red-O staining of formalin fixed cells for detection of lipid accumulation as described previously.^13;16;17^

### RNA-sequencing of iMSCs during chondrogenic differentiation

To examine the transcriptional changes during chondrogenic differentiation using the pellet method, we performed RNA-sequencing of AC- and OA-iMSCs during the course of differentiation process. Pellets from both AC- and OA-iMSCs were isolated at 0, 7, 14 and 21 days of chondrogenic differentiation and total RNA was isolated using miRNeasy Kits. On-Column DNase digestion was performed to remove genomic DNA contamination. RNA quality was checked using Nanodrop and the RNA integrity was determined by Agilent 2200 Bioanalyzer, and the RNA integrity Number (RIN) values were >7 for all samples. Libraries were prepared from 250 ng RNA using TruSeq Stranded Total RNA Sample Prep Kit (Illumina) using the Poly A enrichment method. Sequencing was carried out using the NovaSeq PE 150 system (Novogene UC Davis Sequencing Center, Novogene Corporation Inc.). Raw data were exported in FASTQ (fq) format and quality control was performed for error rate and GC content distribution, and data filtering was performed to remove low quality reads or reads with adaptors. The clean reads were mapped to human reference genome (GRCh38) and differential gene expression (DEGs) analysis was performed using DESeq2 method and pairwise gene expression levels were calculated using RPKM (Read per kilobase of transcript sequence per millions base pairs sequenced) value. FC (Fold Change) in gene expression was performed on filtered data sets using normalized signal values.

### Differential gene expression analysis of RNA-Seq data

Differentially expressed genes (DEG) were identified using DESeq2 in R Bioconductor.^23^ Log fold change (FC) represented the fold change of gene expression, and P<.05 and log_2_FC>2 was set for statistically significant DEGs. Multiple correction testing was performed using False Discovery Rate (FDR). The DEGs between AC- and OA-iMSCs at Day 21 were visualized using heatmap, volcano plot and principal component analysis (PCA) using R-Bioconductor package as described previously.^15;24–26^ Molecular pathways enriched in DEGs was performed using GO (Gene Ontology) and KEGG pathways analysis using STRING (v11.0).^27^ The enrichment of top GO terms based on FDR corrected p-value was visualized by dot plot analysis as described previously.^25;26^ X-axis in the dot plot represents ‘gene term ratio’, which was calculated by ratio of gene numbers enriched in a particular GO term to all the gene numbers annotated in that GO term.

We also performed differential gene expression analysis between healthy and OA chondrocytes by analyzing the publicly available RNA-seq datasets. The raw data were downloaded from healthy and OA cartilage tissues (GSE114007) available from the NCBI-GEO database.^28^ DEGs were identified using DESeq2 in R Bioconductor as described above. The heatmap for mRNA expression profiling of selected genes was generated by R package of pheatmap as described previously.^26^

### Interaction network analysis of DEGs between AC- and OA-iMSCs

To identify the interactions among top DEGs between AC- and OA-iMSCs during chondrogenic differentiation, we performed interaction network analyses using STRING database (v11.0) using a stringent criterion with a combined score of >0.7 showing most significant interactions.^27^ Network clusters were identified using connectivity degree and hub proteins were identified as node showing maximum clustering score in the interaction network. The interaction network was visualized by the Cytoscape (v3.9.0), a bioinformatics package for biological network visualization and data integration^29^ as described previously.^25;26^ Significant clusters in the interaction network were analyzed by sub-network analysis using the Molecular Complex Detection Algorithm (MCODE) plugin (v1.5.1) in Cytoscape.^30^ Enrichment of molecular pathways in identified network cluster was analyzed using ClueGO analysis in Cytoscape.^31^ The genes identified in metabolic and epigenetic regulator pathways in network clusters were also analyzed for differential expression analysis between AC- and OA-iMSCs and visualized by heatmap analysis.

### Statistics

Data are expressed as mean±SEM of at least three independent experiments. All experiments represent biological replicates and were repeated at least three times, unless otherwise stated. Technical replicates are repeat tests of the same value, i.e., testing same samples in triplicate for qPCR. Biological replicates are samples derived from separate sources, such as different clones of iPSCs and iMSCs. Statistical comparisons between two groups (AC- vs OA-) were performed using a two-tailed Student’s t-test for comparing two groups using GraphPad Prism. One-way ANOVA followed by Tukey’s test multiple comparisons test for greater than two groups Significance was denoted at P < 0.05.

---

**Supplementary Figure 1:**
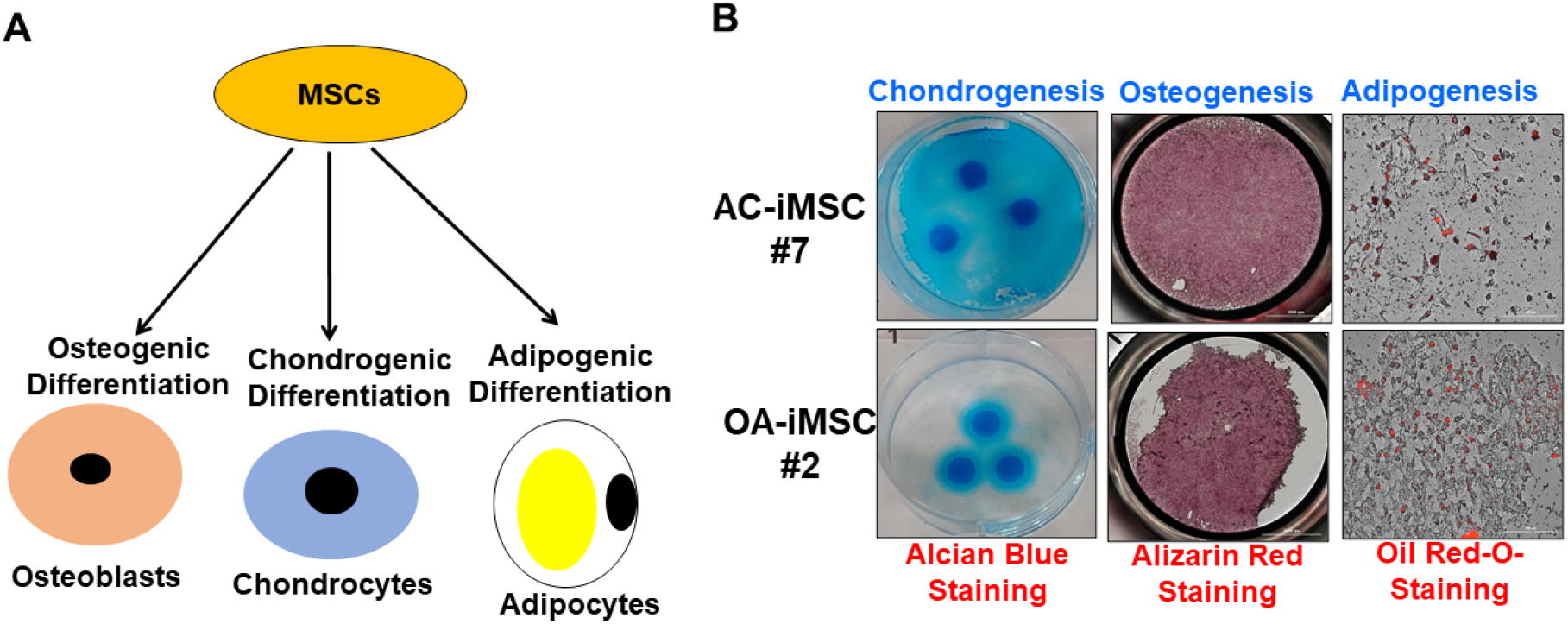
**A)** Trilineage differentiation of these AC- and OA-iMSCs using *in vitro* adipogenic, osteogenic, and chondrogenic differentiation assays. (**B**) Chondrocyte differentiation of AC- and OA-iMSCs was shown by Alcian blue staining of micromass cultures. Osteoblast differentiation was shown by Alizarin Red staining and Adipogenic differentiation was visualized using Oil Red O staining at day 21 culture. Scale bar, 100 μm.

**Supplementary Figure 2:**
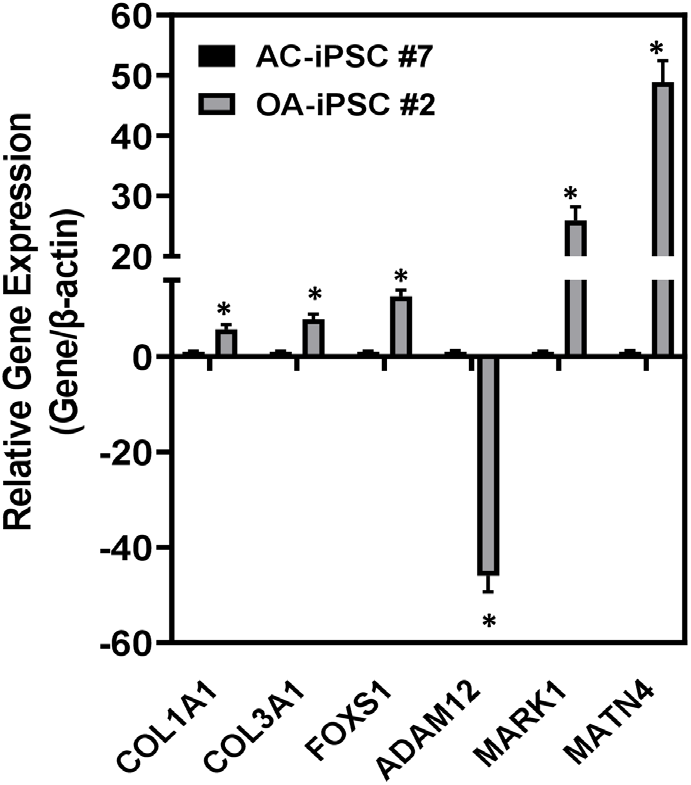
Validation of RNA-Seq results by quantitative PCR. Select genes were chosen for further analysis and validation of the RNA-Seq results. RNAs were isolated from day 21 chondrogenic differentiation of AC- and OA-iMSCs.

## AUTHOR CONTRIBUTIONS

H.D., R.M.G. and N.M.K. conceived the study and designed the project; N.M.K., M.E.D.H. S.C. P.P. performed the experiments; R.M.G. generated the iPSCs; N.M.K. drafted the manuscript; H.D., P.B., and R.M.G. critically reviewed the manuscript. All authors provided the final approval for this manuscript.

## ACKNOWLEDGEMENTS

This work was supported by Georgia and funds from Veteran Affairs and Emory University School of Medicine.

## DATA AVAILABILITY

All raw data has been made available as source data files within the manuscript. The sequencing datasets are available via the Gene Expression Omnibus (GEO) under the accession numbers GSE 214987.

Nazir M Khan and Hicham Drissi, 2022, NCBI Gene Expression Omnibus ID GSE214987. RNA-seq during chondrocyte differentiation of iMSCs derived from iPSCs of healthy (AC-iPSCs) and OA chondrocytes (OA-iPSCs) https://www.ncbi.nlm.nih.gov/geo/query/acc.cgi?acc=GSE214987

